# Wax-printing-free fabrication of paper-supported 3D cancer cell culture

**DOI:** 10.64898/2026.02.10.705119

**Authors:** Ankit Kumar, Bhushan J. Toley

**Affiliations:** Department of Chemical Engineering, Indian Institute of Science, Bangalore, Karnataka, 560012; Department of Bioengineering, Indian Institute of Science, Bengaluru, Karnataka 560012, India

**Author notes:** <u>Correspondence to:</u> Bhushan J. Toley, Department of Chemical Engineering, Indian Institute of Science, Bangalore, Malleswaram, Bangalore 560012.

**Keywords:** tumor microenvironment, hypoxia, doxorubicin, paclitaxel, drug diffusion, drug screening

## Abstract

Three-dimensional (3D) in vitro tumor models are critical for studying transport-limited drug efficacy in solid tumors; however, many existing platforms are technically complex and remain difficult to access. Stacked paper-based tumor models (“cells-in-gel-in-paper”, CiGiP) address this challenge by enabling formation of diffusion-limited microenvironments while allowing direct access to cells from distinct tissue depths. Nevertheless, current CiGiP implementations rely on wax or hydrophobic barrier patterning of paper, which has become increasingly inaccessible. Here, we present a wax-printing-free approach for fabricating stacked paper-supported 3D tumor tissues using a simple 3D-printed press-fit enclosure that holds circular paper layers snugly together, thereby enforcing one-dimensional transport without lateral leakage. Using MDA-MB-231 breast cancer cells embedded in Matrigel, we demonstrate the formation of nutrient-limited microenvironments across the tissue depth, as evidenced by layer-dependent cell viability. The platform enables direct quantification of spatial and temporal drug responses, demonstrated using doxorubicin and paclitaxel, both individually and in combination. Layer-dependent cytotoxicity was measured, and combination treatment analysis revealed antagonistic interactions consistent with prior reports. By eliminating the need for hydrophobic patterning, this approach substantially lowers the technical barriers to constructing stacked paper tumor models and is expected to facilitate broader adoption of paper-supported 3D tissues for drug screening and mechanistic studies.

**Graphical Abstract:** 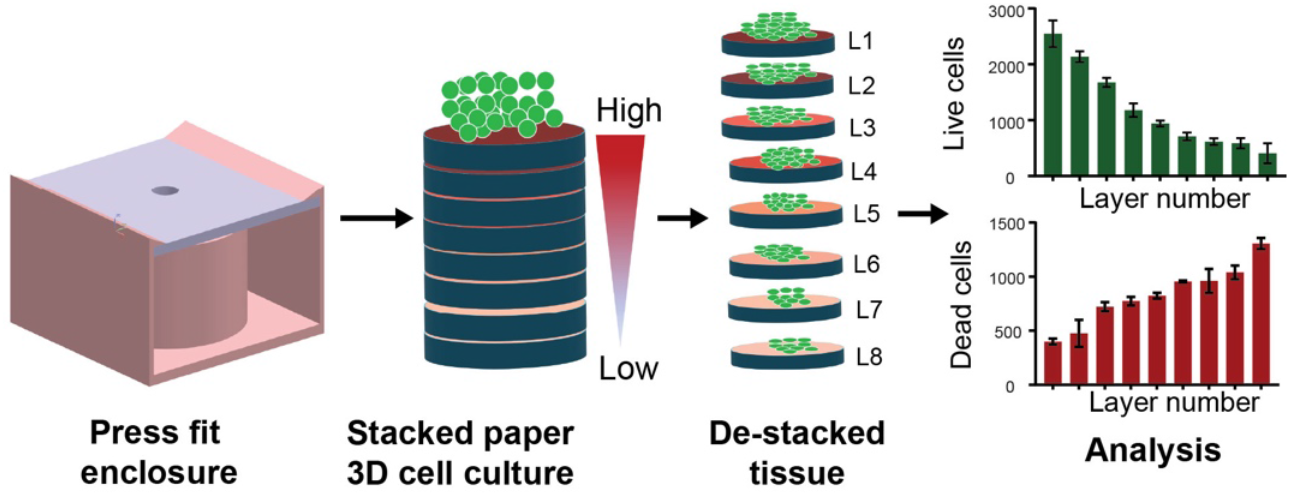

## Introduction

The underdeveloped and chaotic vasculature in solid tumors leads to the formation of oxygen and nutrient gradients, characteristic of the unique tumor microenvironment^1^. These gradients, in turn, lead to the formation of heterogenous regions within the tumor; well vascularized regions contain proliferating cancel cells, while regions far away from a vessel contain quiescent and necrotic cells^1^. Cancer cells in these diverse microenvironments have a diverse response to anticancer agents, with drug transport limitations being an important determinant of drug efficacy^2^. Conventional monolayer cancer cell cultures fail to recapitulate these transport limitations and therefore their utility as an in vitro model of solid tumors is limited.

The significance of in vitro 3D tumor models in cancer research is well understood, and several successful in vitro models have been developed. Of these, cancer spheroids have been one of the most widely used models. Spheroids form by aggregation of cancer cells and feature gradients of oxygen and nutrients, impose transport limitations on drugs, and form heterogenous microenvironments, critical for studying tumor physiology and treatment resistance^3–5^. Scaffold-based or hydrogel-based in vitro 3D tissues, formed by suspending cancer cells in natural or synthetic extracellular matrices, are another widely used in vitro 3D tumor model^6,7^. Recently, 3D bioprinting has emerged as a cutting-edge technique for the fabrication of in vitro 3D tumor constructs^8,9^. Although in vitro 3D tumor models have been tremendously successful in recreating heterogeneous microenvironments, they share a common limitation, in that it remains challenging to readily access cells residing in distinct microenvironments within the 3D tissue for downstream assays.

In vitro 3D cancer tissues formed by stacking paper layers containing cancer cells (“cells-in-gel-in-paper” or CiGiP)^10,11^ overcome the challenge of access to cells from different microenvironments. This model comprises of cancer cells mixed with a compatible extracellular matrix, seeded within layers of paper, which are subsequently stacked. The stack is subjected to cell culture medium from one side, and the other side is kept sealed. Nutrients and oxygen enter the stack from the medium-side and are consumed as they diffuse through the layers of the stack, causing a microenvironment containing limited oxygen and nutrients in deeper layers of the stack. The stacked paper tissue thus mimics a section of a tumor; the top paper layer is representative of a region close to a blood vessel, and the bottom paper layers represent a region of the tumor far from the vessel. The biggest strength of this model is that the tissue can easily by dismantled by de-stacking the layers, and cells from the different microenvironments can be easily accessed. However, the current method for the creation of stacked paper tissues involves patterning each paper layer with wax or wax-like hydrophobic barriers to define the outer boundary of the tissue on each layer. We conducted an extensive literature survey of all reports of stacked paper tissues (Supporting Information S1) and found that, without exception, all reports utilize a technique to pattern a hydrophobic barrier in paper. Although wax printing was once widely used for patterning paper microfluidic devices, the discontinuation of solid-ink printers by Xerox around 2016-2017 has limited its accessibility to many laboratories. Alternative patterning techniques are few and utilized only by niche laboratories focused on developing paper-based microfluidic devices. This has restricted widespread adoption of the powerful paper-based 3D tissue technology.

To overcome this challenge, in this article, we present a method to develop stacked-paper tissues without patterning paper with hydrophobic barriers. This is accomplished by a 3D-printed press-fit kind of a device in which circular paper layers (containing cancer cells and extracellular matrix) are held together snugly, preventing transport of oxygen or nutrients down the side of the stack. Using live-dead staining, we demonstrate that the number of live cells decreases, and the number of dead cells increases from the top layer to the bottom layer, confirming the creation of diffusion-limited zones deep in the tissue. We further demonstrate how this device may be used to measure live-dead cell counts in the various layers of the tissue after treatment with doxorubicin, paclitaxel, and their combination at various concentrations for various periods of time. By obviating the need for patterning hydrophobic barriers in paper, we anticipate that this method will lower the technical barriers faced by researchers in the creation of stacked-paper tissues and lead to widespread adoption of this technique.

## Materials and Methods

### Cell culture device fabrication

Whatman filter paper grade 1 (Cytiva; Cat No-001-150) with a pore size of 11 microns and a thickness of 200 microns was used as the paper scaffold for cell culture. The paper was cut to desired dimensions using a 50 W CO_2_ laser cutter (VLS 3.60; Universal Laser Systems, Scottsdale, AZ). Each cut paper piece was autoclaved before the introduction of cells and extracellular matrix. The housing of the device was 3D printed using tough PLA (Polylactic acid) in an Ultimaker S3 FDM 3D printer.

### Cell culture conditions

MDA-MB-231 cells were obtained as a kind gift from Prof. Ramray Bhat (Indian Institute of Science, Bangalore). These cells were cultured in DMEM supplemented with 10% FBS (Sigma-Aldrich; Catalog No. F0926), containing 1% antibiotic-antimycotic (Sigma-Aldrich; Catalog No. 890890P) on T25 flasks. For culturing cells in Matrigel in paper, 5 μL cell suspension (10^6^ cells/μL) was mixed with 5 μL Matrigel, and the mixture was introduced into a single paper disc. These papers discs were placed in a 12-well plate (one disc per well), which was then placed in a CO_2_ incubator at 37°C for 3 minutes. Following this, 2 mL culture medium was added into each well, followed by incubation for 24 hours. After 24 hours, the discs containing cells-in-Matrigel were ready for stacking in the culture device.

### Quantification of live and dead cells within paper layers

Calcein AM and Propidium Iodide were used for live and dead cell staining, respectively^12^. Stock solutions were prepared in PBS as 3 µM Calcein AM and 10 µg/mL Propidium Iodide. A 250 μL staining solution was prepared by mixing 5 µL Propidium Iodide, 0.7 µL Calcein AM, and 244.3 µL PBS. Staining was conducted on single paper layers placed in individual wells of a 12-well plate. For stacked paper tissues, the layers were de-stacked before staining and imaging. Culture medium was removed from each well and replaced by 250 μL staining solution followed by 30 min incubation at room temperature. The staining solution was then removed, followed by a PBS wash, followed by the addition of 250 μL PBS in each well, at which point the paper layers were ready for imaging.

Fluorescence imaging was conducted using a Zeiss Axiovert A1 FL-LED inverted fluorescence microscope, equipped with a 10X A-Plan objective. The paper disc in the well was flipped and imaged from the bottom. Multiple fields of view spread across the area of the disc were imaged to account for spatial heterogeneity. Images were analyzed using ImageJ. For obtaining cell count of live or dead cells, images were first converted to a binary format using thresholding. The ‘Watershed’ algorithm was applied to separate touching or overlapping cell, and the number of cells was quantified using the ‘Analyze Particles’ function. For experiments in which fluorescence intensity was considered a surrogate measure of live cell count, the mean gray value (pixel intensity) was measured using the ‘Measure’ function. For background subtraction, the background intensity was measured in cell-free regions and subtracted from the mean value over the entire field of view.

### Drug efficacy assessment studies

Doxorubicin (≥95% purity, HPLC; Catalog No. D1515-10MG) and Paclitaxel (≥95% purity, HPLC; Catalog No. T7402-5MG) were purchased from Sigma-Aldrich (St. Louis, MO, USA). Both drugs were dissolved in DMSO to prepare a stock solution and stored at -20°C. Working concentrations were freshly prepared by diluting the stock solutions(1mg/ml) in culture medium before each experiment. Studies were conducted both on cells in Matrigel in single paper layers and stacked paper layers. Single or stacked layers were treated with doxorubicin (DOX), paclitaxel (PTX), or a combination of DOX and PTX at varying for varying periods of time. Cell viability was evaluated using calcein-AM staining, followed by fluorescence imaging and ImageJ-based quantification.

## Results and Discussion

### Design of the 3D-printed stacked paper tissue device

The stacked paper tissue device comprises two interlocking components: a bottom part housing a hollow cylindrical chamber that accommodates a stack of eight paper discs (pink part; Fig. 1), and a top part containing a second hollow cylinder that inserts snugly into the bottom chamber (blue part; Fig. 1). A stack of eight paper discs (each ∼200 µm thick) is placed at the base of the bottom cylinder. The top cylinder is engineered to exert gentle, uniform pressure on the paper stack, thereby securing it in position. Cell culture medium is added into the medium reservoir located within the top cylinder through an inlet port on its upper surface, and this reservoir sits directly above the paper stack. Complete device dimensions are provided in Supporting Information S2. Briefly, both the base of the bottom part and the top surface of the top part are 40 mm × 40 mm squares, the fully assembled device has a height of 24 mm, and each paper disc has a diameter of 20 mm. Design files for the device are available for download from the Supporting Information (Bottom Part.stl and Top Part.stl) to facilitate easy reproduction via 3D printing.

**Figure 1.**
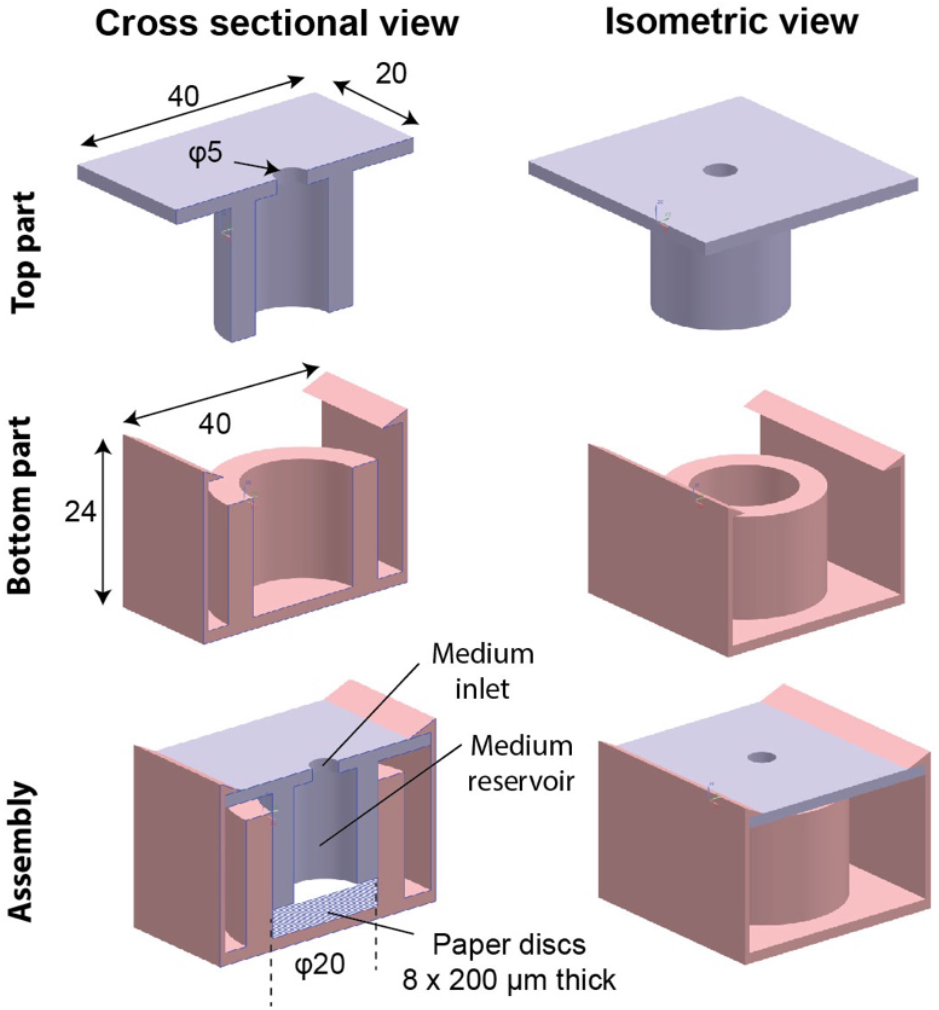
Design of the 3D-printed stacked-paper tissue culture device. All dimensions are in units of mm unless otherwise specified.

### Variation of cell viability across the stacked layers

The choice of membrane and extracellular matrix were important parameters to be established prior to the fabrication of stacked tissues. In preliminary experiments, different paper substrates – Whatman filter paper Grades 1 and 43, glass fiber (Millipore TGFCP203000), Zeiss lens cleaning paper, Whatman lens cleaning paper Grade 105 – were tested for appropriate growth of MDA-MB-231 cells suspended in Matrigel (Supporting Information S3). While there was no significant different in cell proliferation across the membranes, Whatman filter paper Grade 1 was chosen for its mechanical stability and clarity of fluorescence imaging. After selection of Whatman Filter Grade 1 as the support membrane, two different extracellular matrices - Matrigel and Collagen 1 – were compared (Supporting Information S4). There was no significant difference in cell proliferation between the two matrices, and Matrigel was chosen as a more physiologically relevant matrix and for its ability to gel rapidly. As a last experiment to establish the viability of cells in this system, kinetics of MDA-MB-231 cell growth in Matrigel in filter paper membranes was measured (up to 120 h; Supporting Information S5). The number of live cells increased until 72 h of culture, after which it decreased. These experiments validated appropriate growth conditions for cells until at least 72 h of culture in Matrigel in paper.

To create stacked tissues, MDA-MB 231 cells within Matrigel within paper discs, initially cultured in 12-well plates for 24 hours, were arranged and stacked within the device, with growth media supplied from the top. The device was then placed in a CO_2_ incubator at 37°C for 72 hours. Following incubation, the layers were de-stacked, and individual layers were transferred to separate wells of a 12-well plate for live and dead cell staining using calcein-AM and propidium iodide, respectively. A paper layer containing cells in Matrigel that was cultured as a single paper layer in a well plate was used as a reference/control.

Calcein AM staining showed that the number of live cells per field of view (FOV) was the highest in layer 1 (L1) and decreased monotonously from L1 to L8 (Fig. 2A). Similarly, propidium iodide staining showed that the number of dead cells per FOV increased monotonously from L1 to L8. The control layer, which was not nutritionally deprived, had a high number of live cells and a low number of dead cells (Fig. 2A). The number of live and dead cells per FOV in each paper layer were quantified. Compared to the top layer, L1, all lower layers had a lower number of live cells (N=3; **, P<0.01 for L2; and ***, P<0.001 for L3-L8; Fig. 2B), which shows that lower layers of the tissue are nutritionally deficient compared to the top layer. Compared to the control layer, L1 did not have significantly different number of live cells (ns; Fig. 2B), confirming that the top layer receives sufficient nutrition from the medium. Similarly, compared to L1, lower layers L3-L8 had a higher number of dead cells (N=3; *, P<0.05 for L3-L5; and **, P<0.01 for L6-L8; Fig. 2C). Compared to the control layer, L1 did not have significantly different number of dead cells (ns; Fig. 2C). These results demonstrate that the stacked paper tissue successfully recreates a microenvironment analogous to a vascularized tumor region. Layer L1 behaves like tissue close to a blood vessel, receiving adequate nutrients, whereas progressively deeper layers experience increasing nutritional deficiency, mimicking regions of the tumor located farther from the vasculature.

**Figure 2:**
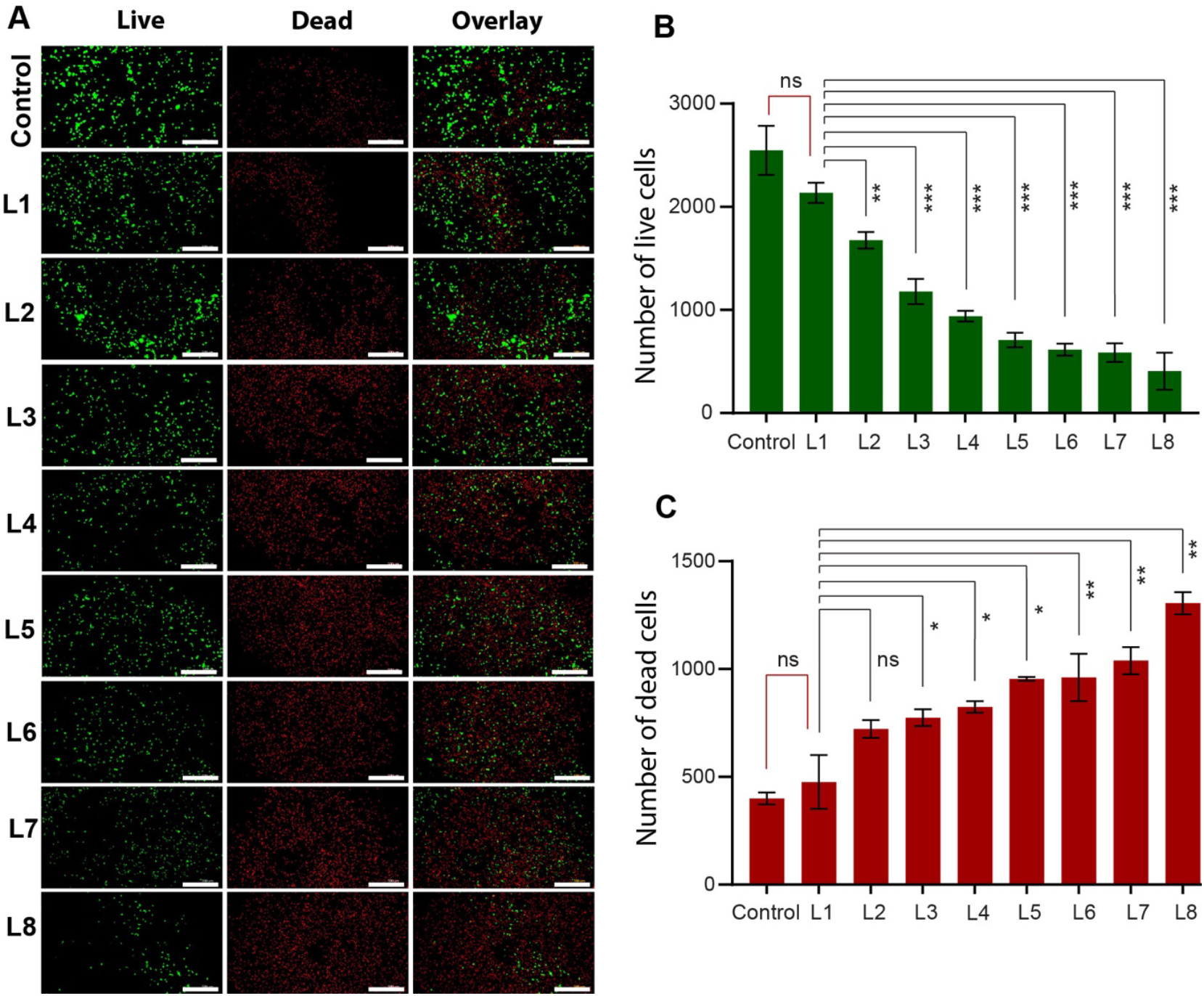
Variation of cell viability across layers of the tissue. **A**. Images of MDA-MB-231 cells in Matrigel in Whatman filter paper Grade 1, stained with Calcein-AM (green; Live), propidium iodide (red; Dead), and merged (Overlay). Control: cells grown in Matrigel on a single paper layer. L1-L8: cells grown in layers 1 to 8, respectively, of the stack. Scale bars are 500 µm. **B-C**. Plot of the number of live (**B**) and dead (**C**) cells per FOV against the layer number (L1-L8). Error bars show standard deviations (N=3 culture devices).

### Doxorubicin (DOX) concentration profile in the stacked paper tissue

The capability of the stacked-paper tissue culture to measure the distribution of a chemotherapeutic agent across the layers is demonstrated next. For this experiment, DOX was chosen as the chemotherapeutic agent because of its inherent fluorescent nature. MDA-MB-231 cells were grown for 48 hours in the stacked-paper tissue culture device. Following this, the culture medium was replaced with medium containing 100 µM DOX, and the treatment continued for 4 hours. Immediately thereafter, the tissue was unstacked and the individual layers were imaged (Fig. 3A). The number of fluorescently labelled cells – reflecting DOX uptake – decreased progressively from the top layer (L1) to the bottom layer (L8) (Fig. 3A). The average fluorescence intensity in each layer, normalized to that of a separately cultured control layer, is shown in Fig. 3B. The mean DOX fluorescence intensity in L1 was not significantly different from the control (ns; Fig. 3B) but was significantly higher than that in the deeper layers (L3–L8; *P* < 0.05; *N* = 3). These results confirm that transport limitations reduce drug availability in the deeper regions of the stacked tissue, thereby recapitulating a transport-limited tumor microenvironment. In another demonstration of the stacked paper device’s capability to mimic transport limitations, the distribution of fluorescent molecules of two different molecular weights – FITC-Dextran 4 kDa and 40 kDA – was also measured (Supporting Information S6). The results confirmed that the larger molecule faced significantly increased resistance for diffusion into the tissue.

**Figure 3:**
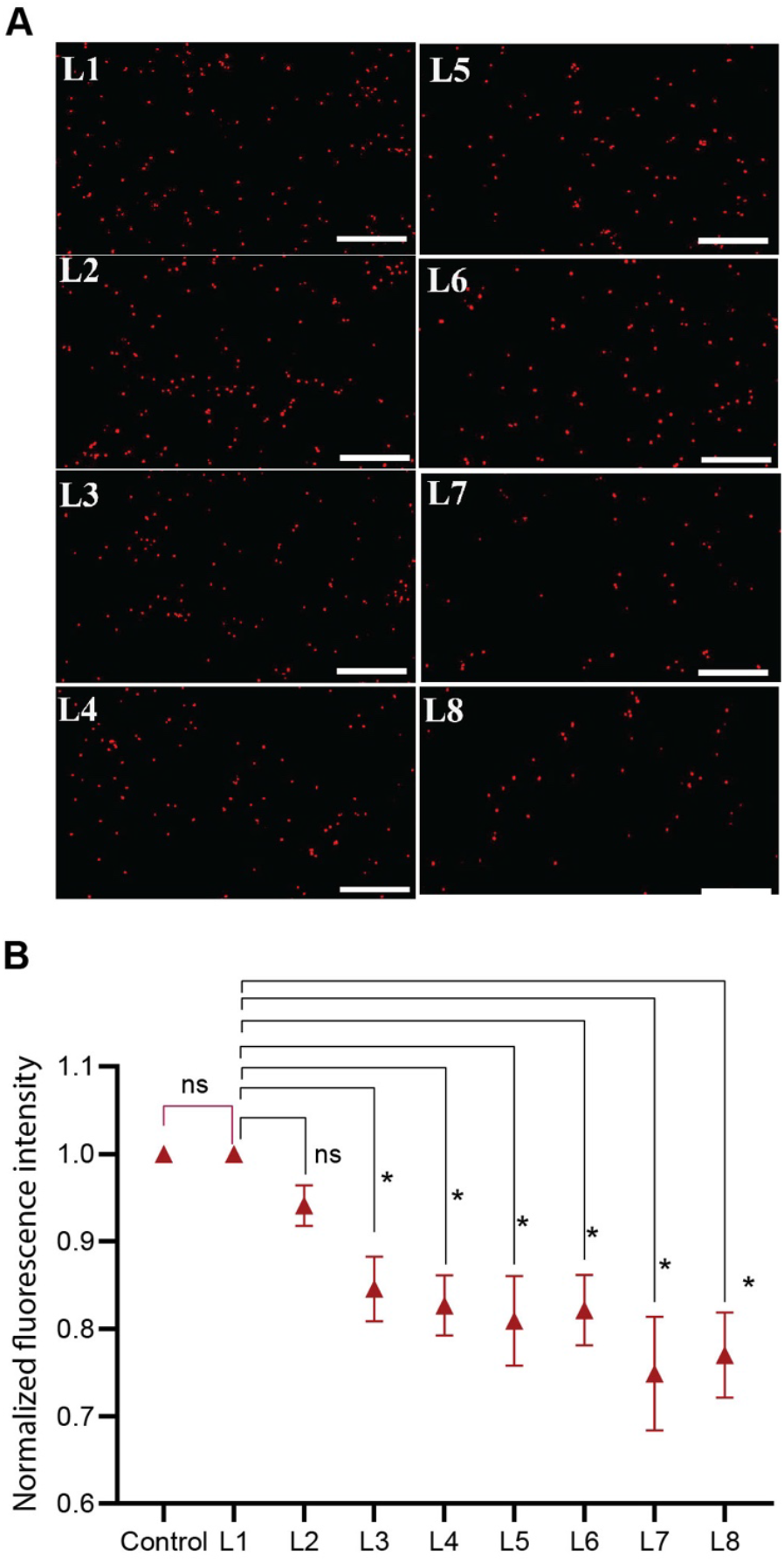
Doxorubicin concentration profile. **A**. Fluorescence images of stacked paper layers illustrating intracellular doxorubicin uptake by MDA-MB-231 cells. Scale bars represent 500 µm. **B**. Plot of doxorubicin fluorescence intensities across stacked paper layers L1-L8, normalized by the fluorescence intensity in a control layer. Error bars show standard deviations (N=3 culture devices).

### Quantification of spatiotemporal response to doxorubicin treatment

The ability of the stacked paper tissue to quantify spatiotemporal drug response is demonstrated next. Stacked paper tissues were formed and cultured in the devices for 24 hours before initiating DOX treatment. In the first set of experiments, all tissues were exposed to 100 µM DOX for varying durations of 4, 8, 16, or 24 h. Regardless of treatment duration, all tissues were de-stacked and analyzed 24 h after the initiation of treatment. For tissues treated for less than 24 h, the DOX-containing medium was replaced with fresh medium after the designated exposure period, and incubation was continued until the 24 h time point, at which the tissues were de-stacked and stained with live-dead cell stains. The number of live and dead cells per FOV as a function of layer number is plotted in Fig. 4A and 4B, respectively. The following trends were observed. The number of live cells across all layers decreased with increasing treatment time (Fig. 4A). For all doxorubicin treatment times, layers L1-L3 had fewer live cells compared to deeper layers, indicating that transport limitations reduced drug efficacy in deeper tissue layers (Fig. 4A). For longer treatment times of 16 and 24 h, the variation in the number of live cells across the layers was less compared to shorter treatment times of 4 and 8 h (Fig. 4A), as diffusion limitations were partially overcome during the longer treatment. A short treatment time of 4 h exhibited an intermediate tissue region (L4-L5) with maximum cell viability (green line; Fig. 4A), which could be indicative of a resistant phenotype developing in these intermediate layers. For the number of dead cells, all these trends were reversed, as expected (Fig. 4B).

**Figure 4:**
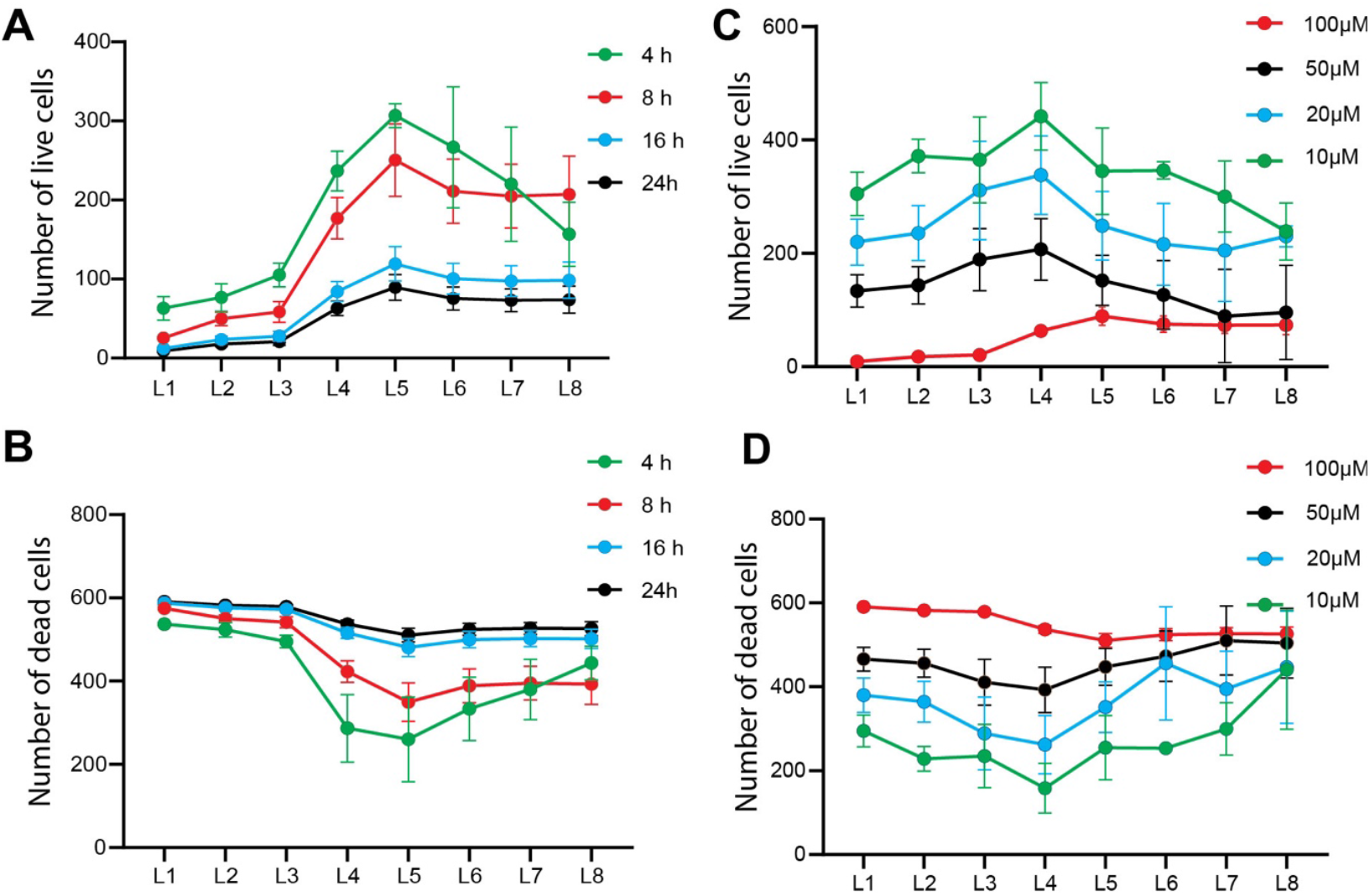
Quantification of doxorubicin-induced cytotoxicity. **A-B**. Number of live (A) and dead (B) cells per FOV, as a function of layer number, after treatment with 100 μM doxorubicin for varying durations. **C-D**. Number of live (C) and dead (D) cells per FOV, as a function of layer number, after treatment with varying doxorubicin concentrations for a fixed duration of 24 hours. Error bars show standard deviations (N=3 culture devices).

In the second set of experiments, tissues were treated with varying DOX concentrations of 10, 20, 50, and 100 μM, but for a fixed treatment time of 24 h. Tissues were de-stacked for analysis immediately after the 24 h treatment. The following trends were observed. The number of live cells across all layers decreased with increasing DOX concentration (Fig. 4C). Because of the long treatment time of 24 h, which provides sufficient time to overcome diffusion limitations, the variation in the number of live cells across the layers was less for all doxorubicin concentrations (Fig. 4C). The number of dead cells exhibited the reverse trend with the number of dead cells increasing with increasing DOX concentration (Fig. 4D).

### Quantification of response to combination treatments

The capability of the stacked paper device to quantify the effect of drug combinations in the context of a transport-limited microenvironment was demonstrated next. Preliminary experiments were conducted on cells grown in Matrigel in single paper layers, which were treated with varying concentrations of DOX and paclitaxel (PTX) (Supporting Information S7). After methods were established, similar experiments were conducted with stacked paper tissues. The stacked constructs were treated with 25µM DOX, 25µM PTX, and a combination DOX+PTX, 25µM each, for 72 hours. After treatment, tissues were de-stacked and the cell viability in each layer was measured. Control tissues, not treated with any drug, were also included. For controls, there was a monotonous reduction in the number of live cells from the top layer, L1, to the bottom, L8 (Fig. 5A), matching the trend in Figure 2B. The effect of the three different treatments – DOX, PTX, and DOX+PTX – on cell viability is shown in Figure 5B. The trend of viability as a function of layer number was consistent across the three treatments. Viability was lowest in the topmost layers that experienced the highest concentration of drug and increased with increasing layer number until L6 (Fig. 5B). Viability decreased in L7 and L8, which is likely due to nutrient deprivation in these layers. Importantly, cell viability in all layers of the treated tissue was less compared to even the deepest layer, L8, of the control tissues. This suggests that drugs were effective across all layers, but intermediate tissue layers may be resistant to drug action.

**Figure 5:**
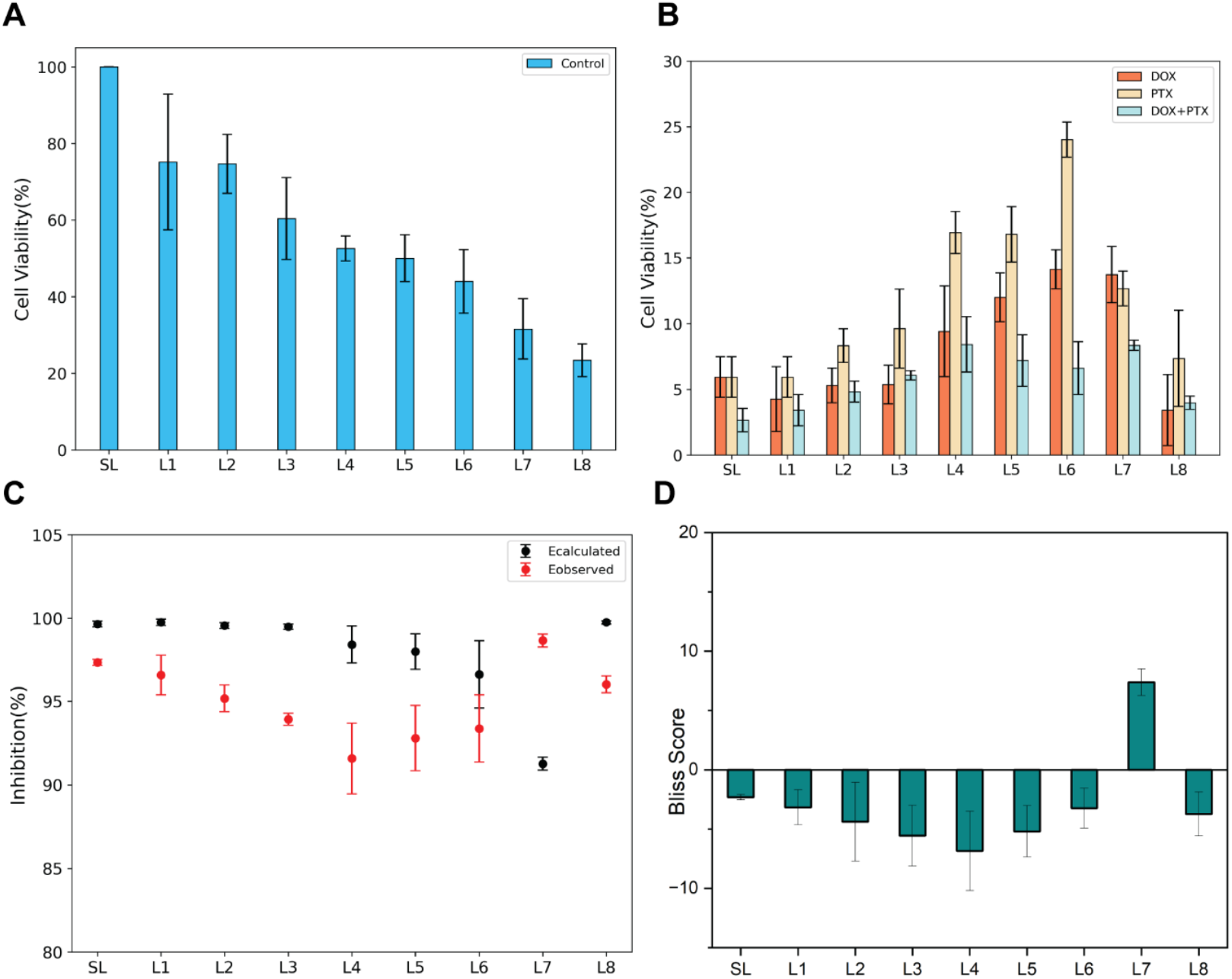
Quantification of the effects of DOX-PTX combination therapy. **A**. Percent cell viability across the stacked layers for control tissues not treated with drugs. **B**. Percent cell viability across the stacked layers for tissues treated with DOX, PTX, or a combination of DOX-PTX. **C**. Observed cell growth inhibition observed experimentally (red markers) and calculated using Bliss independence (black markers). **D**. Bliss score for each paper layer, indicative of drug synergy (positive) or antagonism (negative). Error bars show standard deviations (N=3 culture devices).

To assess whether the combination of drug treatments is synergistic or antagonistic, the Bliss synergy score was calculated^13^. According to this method, first the inhibitory effect, *E*, of each treatment is calculated as the percentage of cells inhibited, or 100 – percentage of live cells. Experimental data is thus used to calculate *E*_*DOX*_, *E*_*PDX*_, and *E*_*DOX*+*PDX*_. The expected combined effect of the drugs, assuming that the drugs act independently is:

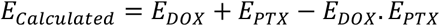

This calculated effect is then compared to the experimentally observed effect, *E*_*observed*_ = *E*_*DOX*+*PDX*_. The Bliss score is then calculated as:

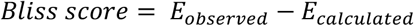

When this score exceeds zero, then the actual effect of the drug combination, *E*_*observed*_, is greater than the expected outcome, *E*_*bliss*_, assuming the drugs act independently. Thus, a positive score indicates synergy, and a negative score indicates antagonism. For the stacked tissue, *E*_*observed*_ and *E*_*calculated*_ for the 8 paper layers is plotted in Fig. 5C. The resulting Bliss score for all layers is plotted in Figure 5D. For all layers (except L7), the Bliss score was negative, showing that DOX and PTX have an antagonistic effect on each other. Such an antagonistic effect of simultaneous DOX+PTX treatment has previously been reported^14^. The anomalous value observed for L7 is most likely attributable to experimental measurement variability.

## Conclusion

This research presents a simple 3D-printed press-fit enclosure that enables consistent assembly of stacked paper-supported 3D tissues. This is the first report of stacked paper-supported tissues without using wax/hydrophobic patterning of paper for defining the culture area. One-dimensional nutrient and drug gradients are ensured by appropriate design of the enclosure, providing a snug fit that prevents transport of drugs and nutrients around the edges of the paper layers. Paper layers can easily be de-stacked by opening the enclosure. We have demonstrated the device’s capability to measure cell viability across the layers in response to drug treatment. The enclosure can be printed on basic FDM 3D printers that are now readily available at a very low cost. The design files are provided for downloading and easy reproduction of the device. We believe that this technique significantly reduces the barriers faced by researchers in the adoption of stacked paper 3D tissues, which are a powerful tool for tissue-based bioassays.

## Supporting information

Device Designs

## Author Contribution Statement

**AK**: Methodology, Investigation, Formal analysis, Writing – Original Draft; **BJT**: Conceptualization, Methodology, Writing – Review & Editing, Supervision, Funding Acquisition.

## Acknowledgements

This work was supported primarily by a CSR grant from Syngene International Ltd. Additional support was provided by an extramural research grant (BT/PR50679/MED/32/998/2023) from the Department of Biotechnology, India; a core research grant (CRG/2021/003397) from the Science and Engineering Research Board, India; and a research grant by the Isaac Center for Public Health (ICPH) at the Indian Institute of Science.

