## Supplementary Information for "Wax-printing-free fabrication of paper-supported 3D cancer cell culture"

**S1: Literature review**

**Table S1:** Compilation of reports of paper-based tissue engineering

| Study | Stack Size | Cell Type | Detection/Gradient | Measurement Tool | Flow Control |
|---|---|---|---|---|---|
| Derda et al. (2009)[1] | 1–8 layers | MDA-MB-231 | Oxygen/Nutrient | Viability, gene expression | Wax printed barrier |
| Mosadegh et al.[2] (2014) | 3–8 layers | Cardiomyocytes | Oxygen | Immunostaining, viability | Wax barrier (CiGiP) |
| Kenney et al. (2015)[3] | 6–10 layers | MDA-MB-231 | Oxygen | Luminescent thin films | PDMS tape |
| Chung et al. (2018)[4] | ~10 layers | N/A | pH | pH-sensitive films | Adhesive lamination |
| Derda et al.[5] | 8 layers | A549 | Cell migration study | GFP-expressing | Wax barrier |
| Deiss et al[6]. | 3 layers | Human breast cancer cells | Drug screening | N/A | Patterned with wax and PDMS tape |
| Mosadegh et al.[7] | 3 layers | MDA-MB-231 | Cellular invasion/oxygen gradient | fluorescence microscopy | Wax barrier |
| Darren et al.[8] | 6 layers | N/A | oxygen gradient | Fluorescence imaging | Tissue roll |
| Tobias et al.[9] | N/A | N/A | Drug screening | Mass spectroscopy | wax-patterned paper |
| Dermutz et al.[10] | network topology | N/A | intracellular and extracellular signals | Fluorescence imaging | hydrophobic barrier with a solid-ink printer |
| Lei et al.[11] | Paper-based microreactor | N/A | Cellular phosphorylation, cell morphology | Immunoassay | Wax barrier using wax printing |
| Zang et al.[12] | Paper-based electrochemical immunoassa | N/A | N/A | Electrochemical immunoassay | Wax-patterned paper |

| | | | | | |
|---|---|---|---|---|---|
| | y device (3D-Mpeid) | | | | |
| Juvonen et al.[13] | 24-wells array | ARPE-19 Cells | Cell culture | Contact angle, surface energy, AFM | flexographically printed PDMS film |
| Hong et al.[14] | Serpentine channel | N/A | Drug discovery | Concentration gradient for DOX, cell viability | hydrophobic barrier using native photoresist SU-8 3010 |
| Pitaksit et al.[15] | custom-designed 3D-oriented transwell device | Caco-2 | 3D Microenvironment for cell growth, proliferation, and differentiation | Immunofluorescence and SEM analysis, TEER | wax-patterned |
| Rahimi et al.[16] | Stacked paper layer | ALI-cell | New drug discovery | Fluorescent imaging | Direct-patterned laser-treated hydrophobic barrier |
| Zachary et al.[17] | Perfusion-based system | Endothelial cell | Co-culture in the absence of an abiotic gradient, spheroids-on-demand, and blood vessel-like structures | Fluorescent imaging | Wax printed to create a hydrophobic barrier |
| Lee et al.[18] | Patterned based stacked layer | Huh-7 | Drug discovery, cell proliferation | SEM, WST-1 assay | Wax printed barrier-Cellulose paper |
| Smith et al [19] | Stand-alone paper based plateform | HDF and Hela cells | Drug screening, cytocompatibility studies | AFM, Barrier test | Nanostructured barrier |
| Sapp et al.[20] | 4 layers of 114 filter paper | Aortic valves cells | Cell viability and cell migration studies | Gel and confocal microscopy imaging | Wax printed |

**S2. Design of the 3D-printed stacked paper tissue device**

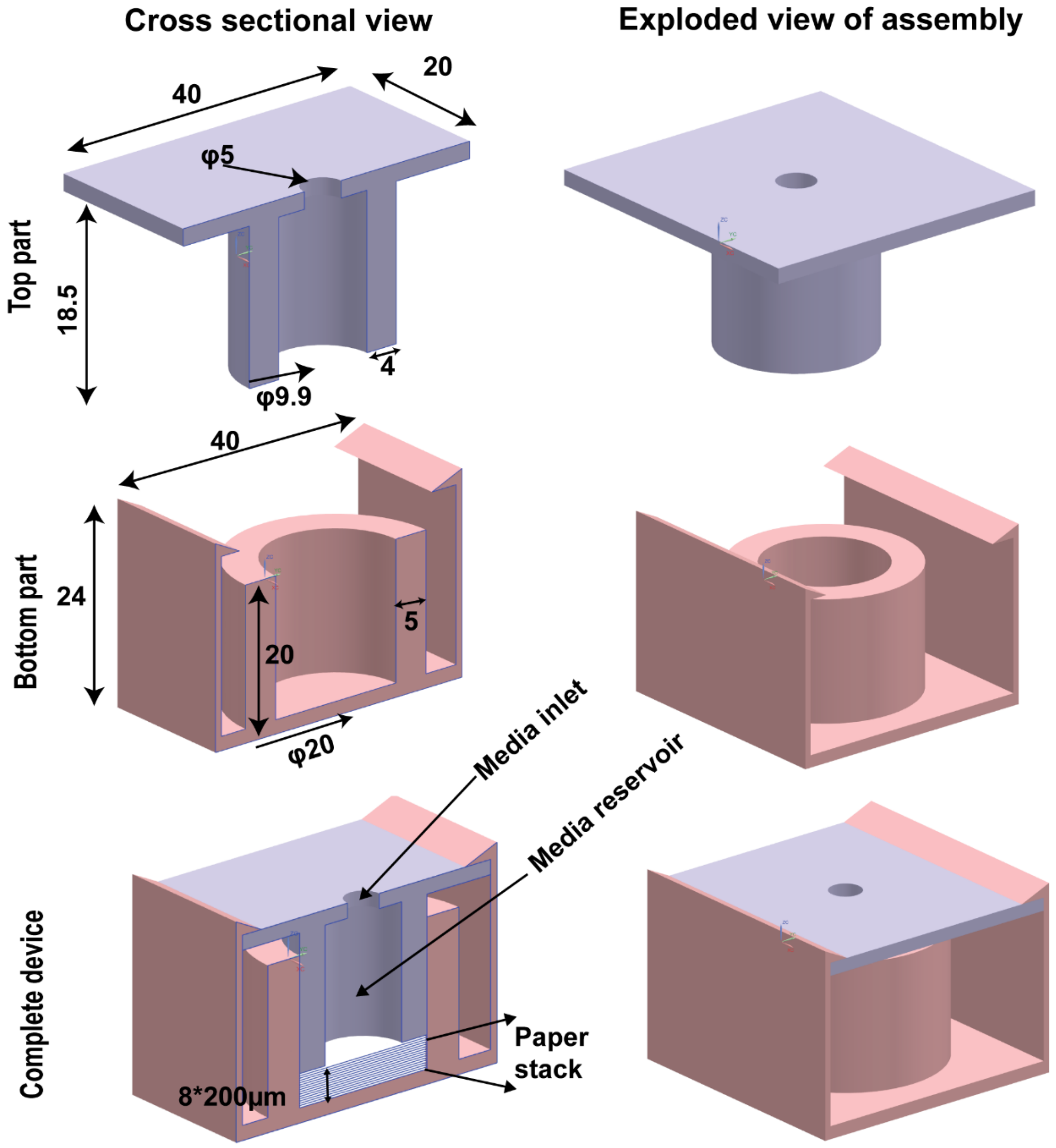


**Figure S2**:Design of the 3D-printed stacked-paper tissue culture device. All dimensions are in units of mm unless otherwise specified.

**S3. Culture of MDA-MB-231 cells on different paper-based substrates in Matrigel**

MDA-MB-231 cells, a model for aggressive breast cancer, are cultured on various substrates to study their growth and behaviour. The percentage of live and dead cells was calculated as:

$$\%\,Live\ Cells = \frac{Number\ of\ live\ cells}{Number\ of\ live\ cells + Number\ of\ dead\ cells} \times 100$$

$$\%\,Dead\ Cells = \frac{Number\ of\ dead\ cells}{Number\ of\ live\ cells + Number\ of\ dead\ cells} \times 100$$

No significant difference in cell proliferation was observed across the membranes. Whatman filter paper 1 was chosen as the membrane to proceed with because of its mechanical stability during destacking and the clarity of fluorescent images generated.

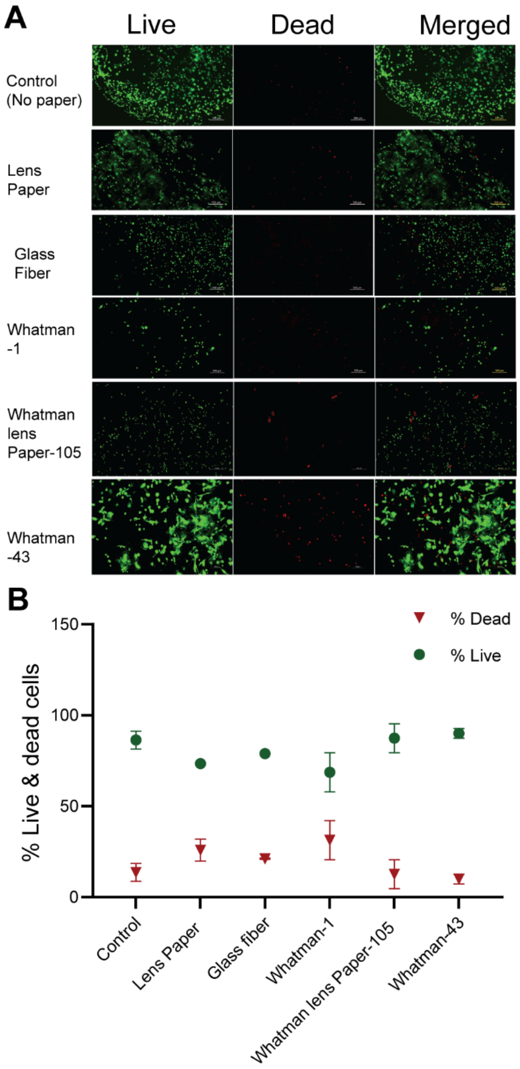


**Figure S3**:(A) Live and dead cell-stained image of MDA-MB-231 cells grown in Matrigel (10x magnification). The scale bar in the image is 200 µm (B). Plot of percentage live and dead cells in different papers.

**S4. Growth and behaviour of MDA-MB-231 cells on various substrates involving collagen and Matrigel**

MDA-MB-231 cells were grown under 4 conditions – i) in collagen without paper, ii) in paper without collagen, iii) in collagen in paper, and iv) in Matrigel in paper. Percentage live cells after a growth period of 48 hours is shown in Figure S4. There was no significant difference between conditions iii) and iv), and cells in Matrigel in paper was chosen as the matrix of choice for its physiological relevance as well as its ability to gel rapidly after introduction into paper.

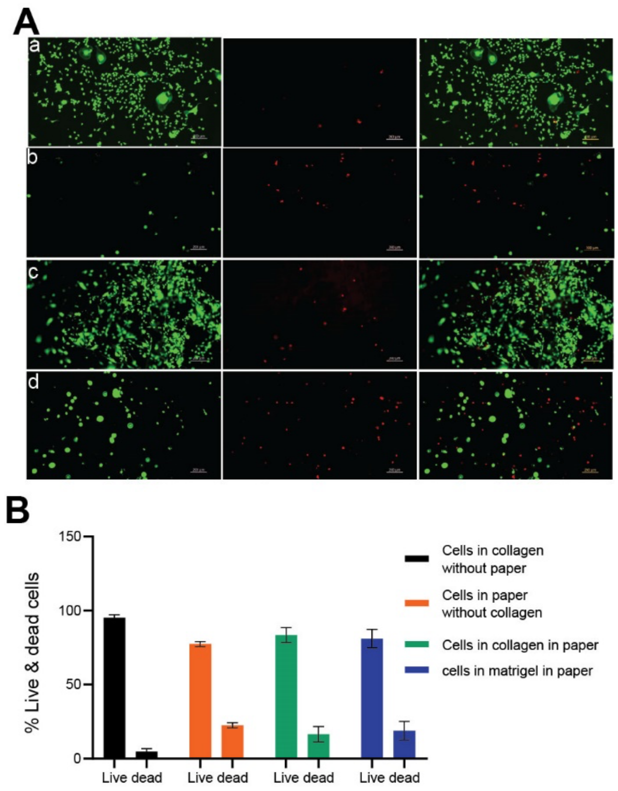


**Figure S4**:(A) Live and dead cell-stained image of MDA-MB-231 cells grown in (a) Cells in gel without paper, (b) Cells in paper without gel, (c) Cells in collagen in paper, (d) Cells in Matrigel in paper at 10x magnification. The scale bar of the image is 200 µm. (B) Plot of percentage live and dead cells in various substrates**.**

**S5. Kinetic study of MDA-MB-231 cells growing in Matrigel in Whatman filter paper Grade-1**

To assess whether MDA-MB-231 cells were proliferating, 0.1 million cells/mL were seeded in Matrigel in Whatman filter paper (Grade 1) placed in 12-well plates. The cells were incubated in an environment consisting of 5% $CO_2$, 37 °C, and 15% humidity for 0, 24, 48, 72, and 120 hours. After each incubation period, the cells were stained with Calcein-AM to label live cells (green fluorescent). The acquired image shows live cells (Figure S5A), and the acquired images were imported into ImageJ and converted to an 8-bit grayscale format. The average pixel intensity was quantified for each image frame using ImageJ for per field of view (FOV) . Intensity measurements were performed for three independent experiments, and the mean values were calculated, as depicted in Figure S5B.

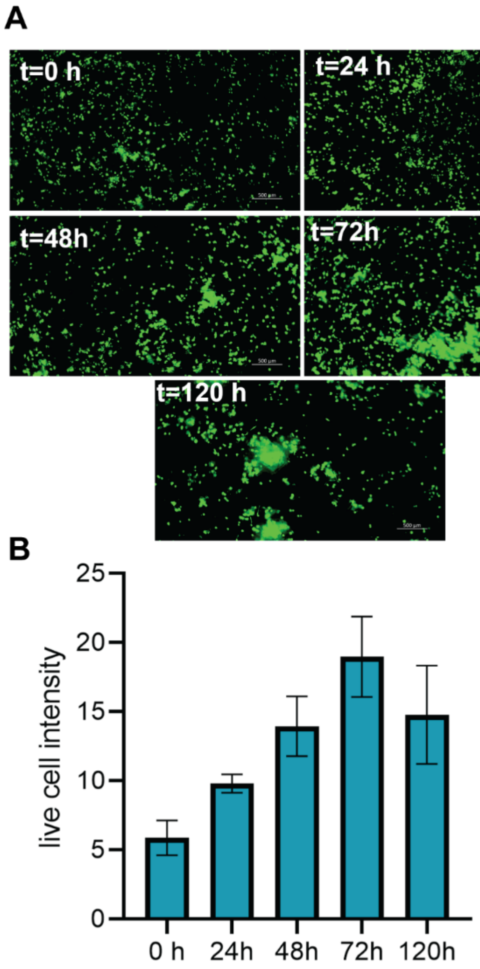


**Figure S5**: (A) Fluorescent image of live(green) MDA-MB-231 cells in Matrigel after staining with Calcein-AM at various times. The scale bar of the image is 200 µm. (B) Plot of live cell intensity at different times.

#### S6. Transport of FITC-dextran of different molecular weights in the 3D paper stack

To investigate the diffusion of molecules with varying molecular weights through stacked paper layers, eight PBS-saturated paper layers were assembled within a 3D-printed device. FITC-dextran – 4 kDa and 40 kDa – was then introduced at concentrations of 100 µM into the top culture chamber. After 1 hour, the layers were de-stacked, and fluorescence images of each layer were captured using a fluorescent microscope. The experiments were performed in triplicate, and the mean and standard deviation of fluorescence intensity were plotted for each layer for both molecules. The intensity of both molecules was similar in L1 and decreased with increasing layer number, but decreased more rapidly for FITC-dextran 40 kDa compared to FITC-dextran 4 kDa (Fig. S6 A-B). This demonstrates the tissue model's ability to accurately reproduce transport limitations faced by molecules of different molecular weights.

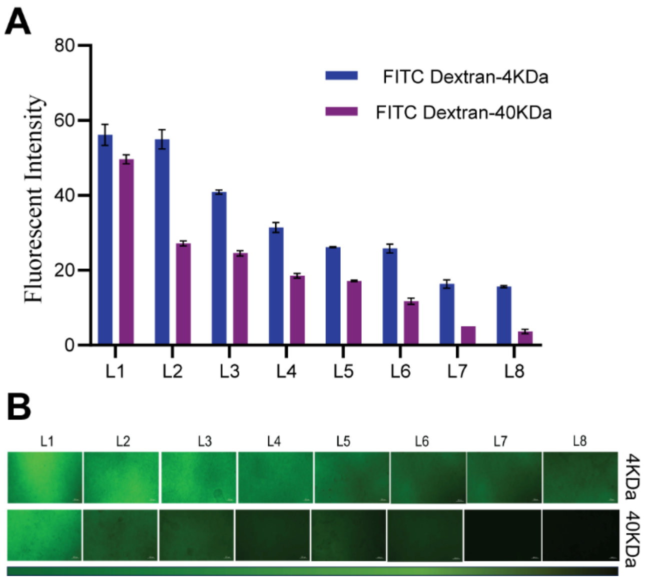


**Figure S6:** (A) Fluorescence intensity profiles of FITC–dextran (4 kDa) and FITC–dextran (40 kDa) across stacked layers of Whatman filter paper grade 1 after 1 h of treatment. (B) Fluorescence images of FITC–dextran (4 and 40 kDa) across stacked layers. The scale bar of the image is 200 µm

**S7. Pharmacodynamic Analysis of Doxorubicin, Paclitaxel, and their Combination in a Single Layer (SL) Cell Culture.**

Cells were seeded at a density of 0.1 million cells/mL in Matrigel in paper, which was then placed in 12-well plates. The cultures were incubated for 48 hours under standard conditions (37 °C, 5% $CO_2$, and a humidified atmosphere) in drug-free medium to allow for equilibration. The medium was then removed, and the cells were washed with PBS. Cells were subsequently treated with doxorubicin (DOX), paclitaxel (PTX), and a combination of DOX and PTX at concentrations of 0.1, 1, 10, 25, and 50 μM. After 72 h of drug exposure, the drug-containing medium was removed, cells were washed with PBS, and cell viability was assessed by staining with calcein-AM. Live-cell imaging was performed using fluorescence microscopy in the FITC channel at 10× magnification. The acquired images were imported into the ImageJ software for quantification of live-cell fluorescence intensity. Images were converted to 8-bit grayscale format, and the average fluorescence intensity per field of view(FOV) was calculated for each image. This analysis was performed for all images and repeated across three independent experimental samples.

In Figure S7A, it can be clearly observed that the intensity of live cells decreases as the drug concentration increases. This indicates a strong pharmacodynamic effect of the drug, both for the individual treatments and their combination. From these intensity values, percent cell viability was calculated and plotted to show the viability across different concentrations of doxorubicin, paclitaxel, and their combination. Percentage inhibition was calculated as 100 minus the percentage cell viability.

$$\%\ Cell\ Viability = \frac{Live\ cells\ intensity(c)}{Live\ cell\ intensity(c{=}0\mu M)}$$

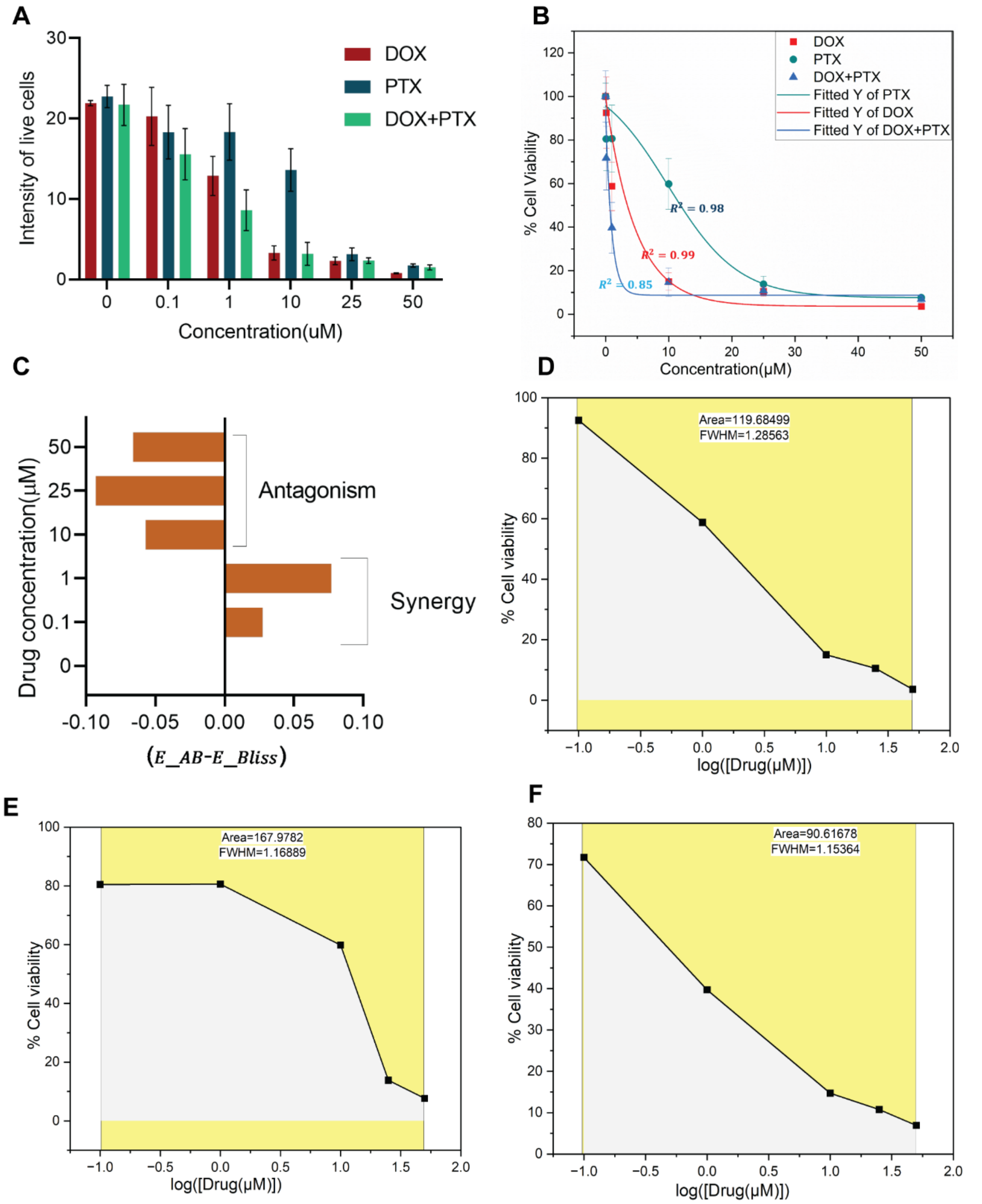


**Figure S7: (A)** Live-Cell Fluorescence Intensity of Cells Treated with DOX, PTX, and DOX+PTX at Varying Concentrations. (B) Dose-Dependent Cell Viability Following DOX, PTX, and DOX+PTX Treatment. (C) BLISS Score Analysis of DOX–PTX Combination Across Drug Concentrations. (D–F) Dose–response curves showing percentage cell viability as a function of log(drug concentration) for DOX (D), PTX (E), and DOX+PTX (F).

Next, we fitted the experimentally obtained percentage viability data to a pharmacodynamic dose–response model (equation S1) to determine the $IC_{50}$ value—a well-known parameter that reflects the potency of a drug (Figure S7(B)). The combination treatment exhibited a lower $IC_{50}$ value compared to the individual drugs, indicating greater potency in the combined formulation (Table S1).

$$Y(\%\ Cell\ Viability) = A_1 + \frac{A_1 - A_2}{1 + 10^{(logIC_{50} - X)\times n}} \qquad (S1)$$

**Table S1:** $IC_{50}$(μM) for all drug treatments in a single layer(SL).

| **Drugs** | **$IC_{50}$(μM)** |
|---|---|
| Doxorubicin(DOX) | 2.24 |
| Paclitaxel (PTX) | 12.1 |
| DOX+PTX | 0.6 |

The BLISS score (methods described in the main text) was then calculated and is reported in Fig. S7C. Finally, the area under the curve (AUC) was determined for the dose–response curves of both individual and combination treatments to assess drug efficacy (Figures S7-D–F). The combination treatment exhibited a smaller AUC compared to the individual drugs, indicating enhanced potency and efficacy in the single-layer (SL) cell culture model.

### References

(1) Derda, R.; Laromaine, A.; Mammoto, A.; Tang, S. K. Y.; Mammoto, T.; Ingber, D. E.; Whitesides, G. M. *Paper-Supported 3D Cell Culture for Tissue-Based Bioassays*. www.pnas.org/cgi/content/full/.

(2) Mosadegh, B.; Lockett, M. R.; Minn, K. T.; Simon, K. A.; Gilbert, K.; Hillier, S.; Newsome, D.; Li, H.; Hall, A. B.; Boucher, D. M.; Eustace, B. K.; Whitesides, G. M. A

Paper-Based Invasion Assay: Assessing Chemotaxis of Cancer Cells in Gradients of Oxygen. *Biomaterials* 2015, *52* (1), 262–271. https://doi.org/10.1016/j.biomaterials.2015.02.012.
